# Single nucleus and spatially resolved intra-tumor subtype heterogeneity in bladder cancer

**DOI:** 10.1101/2022.10.27.513983

**Authors:** Sia V. Lindskrog, Sofie S. Schmøkel, Iver Nordentoft, Philippe Lamy, Michael Knudsen, Frederik Prip, Trine Strandgaard, Jørgen Bjerggaard Jensen, Lars Dyrskjøt

## Abstract

Current transcriptomic classification systems for bladder cancer do not consider the level of intra-tumor subtype heterogeneity. Here we present an investigation of the extent and possible clinical impact of intra-tumor heterogeneity across early and more advanced disease stages of bladder cancer. We performed single nucleus RNA-sequencing of 48 bladder tumors and four of these tumors were additionally analyzed using spatial transcriptomics. Total bulk RNA-sequencing and spatial proteomics data were available from the same tumors for comparison, along with detailed clinical follow-up of the patients. We demonstrate that tumors display varying levels of intra-tumor subtype heterogeneity and show that a higher class 2a weight estimated from bulk RNA-sequencing data is associated with worse outcome in patients with molecular high-risk class 2a tumors. Our results indicate that discrete subtype assignments from bulk RNA-sequencing data may lack biological granularity and continuous class scores could improve clinical risk stratification of patients.

**Highlights:**

- Single nucleus RNA-sequencing of tumors from 48 bladder cancer patients.
- Tumors display varying levels of intra-tumor subtype heterogeneity at single nucleus and bulk tumor level.
- The level of subtype heterogeneity could be estimated from both single nucleus and bulk RNA-sequencing data with a high concordance between the two.
- High class 2a weight estimated from bulk RNA-sequencing data is associated with worse outcome in patients with molecular high-risk class 2a tumors.

## Introduction

Transcriptomic subtyping of bladder cancer (BC) using bulk gene expression profiling has been in focus for two decades^1^. A consensus molecular classification of muscle invasive BC (MIBC) has been established^2^ and the European UROMOL consortium has delineated transcriptomic classes of non-muscle invasive BC (NMIBC) reflecting diverse tumor biology and disease aggressiveness^3,4^. However, while these bulk classification systems assign a single subtype to each tumor, expression profiling of multiple tumor regions and immunohistochemistry analyses have previously shown that different subtypes can co-exist within a single tumor^5,6^.

Single cell technologies provide a new opportunity to study tumor ecosystems and tumor heterogeneity at single cell resolution. While some previous studies have focused on the tumor microenvironment of BC^7,8^, studies of heterogeneity within the epithelial compartment are emerging as well^9–12^. Single cell RNA-sequencing (scRNA-seq) require single cell suspensions from fresh tissue isolated directly from surgery, while single nucleus RNA-sequencing (snRNA-seq) makes it possible to investigate frozen tumor tissue, facilitating biobank studies of clinically well-annotated patients. Interestingly, a recent study performed snRNA-seq of frozen tumors from 25 patients with MIBC and identified a N-cadherin 2 expressing epithelial subpopulation with abilities for therapeutic response prediction^12^. However, studies investigating the extent and possible clinical impact of intra-tumor heterogeneity at single cell resolution in larger cohorts of both early and advanced disease stages are still lacking.

Here, we performed snRNA-seq analysis of 59,052 nuclei from 48 bladder tumors to study tumor heterogeneity at single cell resolution. Additionally, data from total bulk RNA-sequencing (RNA-seq), spatial transcriptomics and spatial proteomics was used for comparison. We demonstrated concordance between immune cell infiltration measured by multiplex immunofluorescence (mIF) and snRNA-seq, and we showed that tumors displayed varying levels of intra-tumor subtype heterogeneity. The subtype heterogeneity was also documented by spatial transcriptomics, and we demonstrated that intra-tumor subtype heterogeneity may affect clinical outcome.

## Results

### snRNA-seq of human bladder tumors; data processing and method comparison

In order to perform retrospective studies on samples from clinically well-annotated patients, we performed snRNA-seq of tumors from 48 BC patients using an optimized DroNc-seq protocol^13^ (**Supplementary Table S1**). As we observed more nucleus barcodes than expected for each tumor and too sparse data per nuclei, an extra processing step of the nucleus barcodes was implemented as follows. First, we generated a whitelist of barcodes for each tumor by selecting the *x* number of barcodes with the highest number of genes expressed, where *x* was based on the theoretical number of nuclei analyzed in the laboratory (**Supplementary Table S1**). Second, we corrected remaining non-whitelisted barcodes to a whitelisted barcode if a Hamming distance of 2 (allowing two base substitutions) was present to collapse barcodes originating from the same barcode. Importantly, the barcode processing increased the number of genes and counts per nucleus, which was essential for downstream analysis (**Figure 1A**). We obtained data from 117,653 nuclei after the initial barcode processing, and 59,052 nuclei passed downstream quality control filtering (**Figure 1B**). To investigate the quality of data generated using DroNc-seq, three samples were subsequently analyzed using the 10x Chromium platform. After quality control filtering, 4,431 nuclei with an average of 568 expressed genes per nucleus remained when using the DroNc-seq platform, whereas 5,758 nuclei with an average of 1,649 expressed genes per nucleus remained when using the 10x Chromium platform (**Figure 1C**). For genes with non-zero expression in both the DroNc-seq and 10x Chromium datasets, we observed 193 and 1,054 counts per gene on average, respectively. Consequently, the 10x Chromium platform generated more data per nucleus compared to DroNc-seq but also at a much higher cost. We assigned all nuclei to one of five major cell populations (epithelial, fibroblast, endothelial, lymphoid or myeloid) using data from a previous study of snRNA-seq of human bladder tumors as reference^12^. Nuclei with a delta median value (the difference between the score for the assigned label and the median across all labels for that nuclei) below 0.05 were excluded to remove low-quality cell type assignments. Uniform Manifold Approximation and Projection (UMAP) visualization of the 10x Chromium data revealed three major epithelial clusters separated by patient origin as well as smaller clusters consisting of fibroblasts, endothelial cells and immune cells (**Figure 1D**). In contrast, non-epithelial cells clustered together with the patient-specific epithelial cells for the DroNc-seq data, probably due to the lower number of genes detected per nucleus (**Figure 1E**). To confirm cell type annotations of the more sparse DroNc-seq data, we compared the relative proportions of cell type annotations within each tumor obtained from the two platforms. We found a similar proportion of predicted non-epithelial cells, confirming that the epithelial compartment constitutes the bulk of these tumors when using either of the two platforms (**Figure 1F**). We did observe differences in the fractions of non-epithelial cell types between the platforms which may be caused by tumor heterogeneity (not the same cells analyzed on both platforms) but also differences in number of genes detected per nucleus (**Supplementary Figure S1A**). Although the comparison of methods showed a better performance of the 10x Chromium platform, the DroNc-seq platform allowed us to analyze a larger cohort of human bladder tumors at single cell resolution due to the lower cost of the analysis.

**Figure 1.**
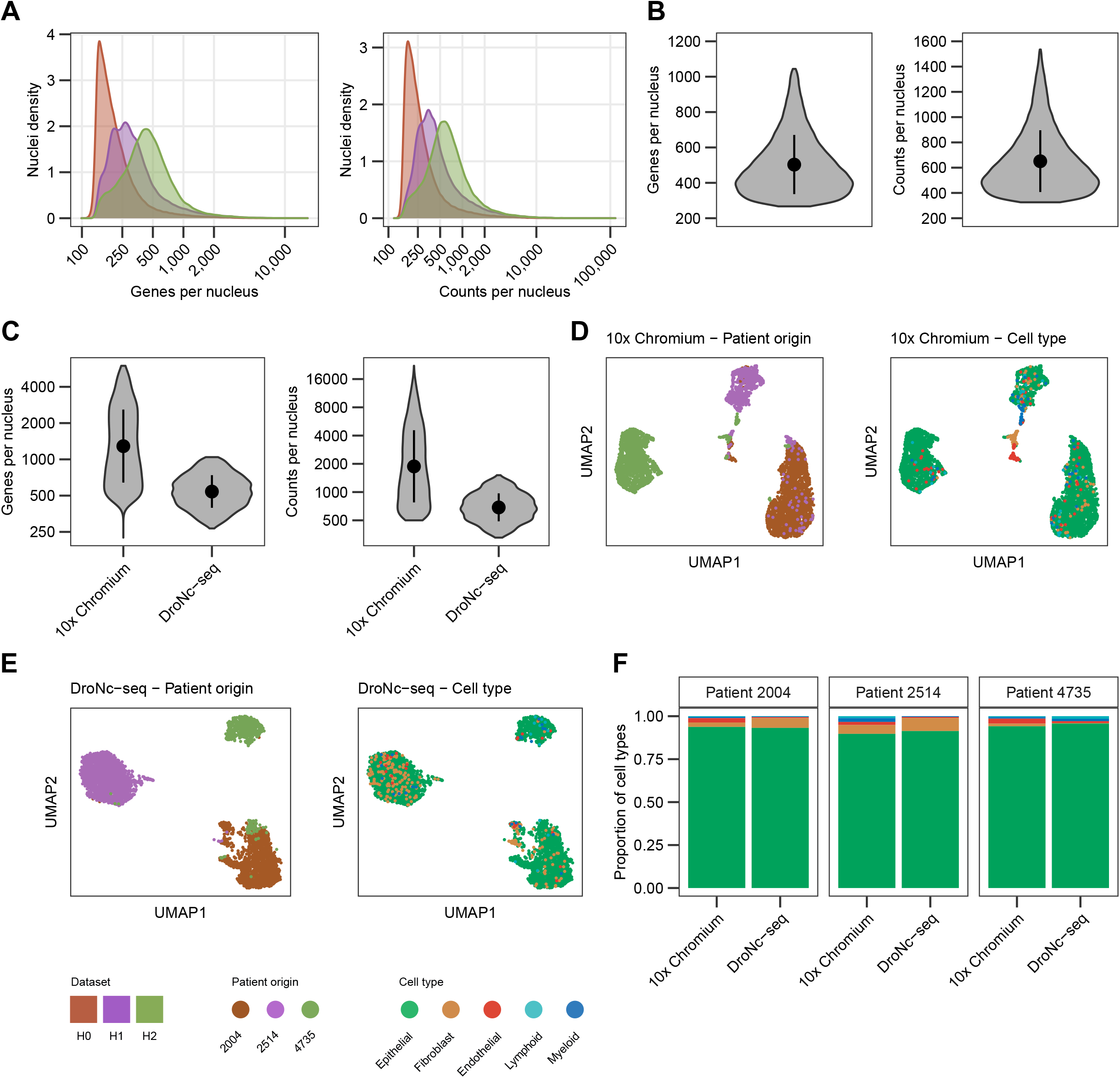
snRNA-seq of human bladder tumors; data processing and method comparison. **A)** Distribution of number of genes (left) and counts (right) detected per nucleus for datasets with different Hamming distances used to correct barcodes. **B)** Number of genes (left) and counts (right) per nucleus for the full DroNc-seq dataset after quality control filtering (n=59,052). **C)** Number of genes and counts per nucleus for three samples analyzed using the 10x Chromium platform (n=5,758) and the DroNc-seq platform (n=4,431). **D)** UMAP visualization of 5,010 nuclei from three samples analyzed using 10x Chromium colored by patient origin (left) and cell type (right). **E)** UMAP visualization of 3,903 nuclei from three samples analyzed using DroNc-seq colored by patient origin (left) and cell type (right). **F)** Relative proportions of cell types within three samples analyzed on two platforms: 10x Chromium (2004, n=2,569; 2514, n=895; 4735, n=1,546) and DroNc-seq (2004, n=1525; 2514, n=1794; 4735, n=584).

### Exploring the tumor microenvironment of human bladder tumors

The DroNc-seq dataset consisted of 48 bladder tumors spanning the disease spectrum of BC (10 Ta, 13 T1, 25 T2-4). In total, 48,567 nuclei were annotated to a major cell type and the various cell populations showed higher expression of corresponding marker genes (**Figure 2A**). Overall, the 48 tumors were predicted to consist of 91% epithelial (cancer) cells, 5% fibroblasts, 3% immune cells and 1% endothelial cells. Graph-based clustering and UMAP visualization of all tumors were mainly driven by patient origin indicating a high level of inter-tumor heterogeneity (**Figure 2B**). The fraction of non-cancer cells varied between tumors and increased with tumor stage, as expected (**Figure 2C**). Other data layers, including mIF and bulk RNA-seq, have previously been generated from the same tumor samples enabling comparison of the estimated percentage of immune cells across these layers^14–16^. We observed that the majority of tumors with a higher amount of immune cells estimated from the DroNc-seq data also had a higher percentage of immune cells estimated from mIF data and a relatively higher immune score estimated from bulk RNA-seq data (**Figure 2D-E**).

**Figure 2.**
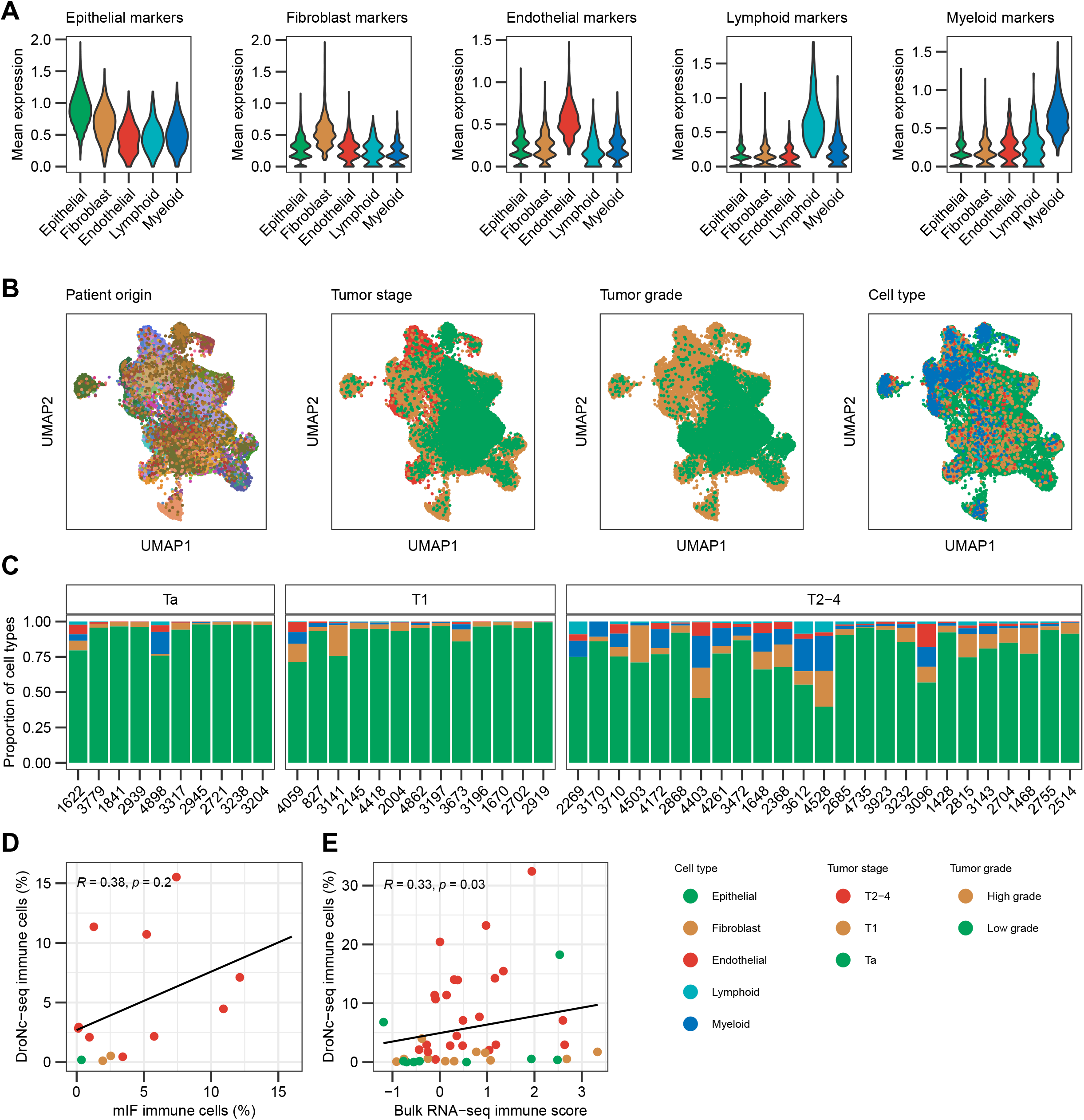
Exploring the tumor microenvironment of human bladder tumors. **A)** Violin plots showing the mean expression of epithelial-, fibroblast-, endothelial-, lymphoid- and myeloid gene markers within the different cell types. Cell type markers were identified from the reference dataset using SingleR. **B)** UMAP visualization of 48,567 nuclei from 48 tumors analyzed using the DroNc-seq platform colored by patient origin, tumor stage, tumor grade or cell type. **C)** Relative proportions of cell types within each sample separated by tumor stage. Samples are sorted by increasing numbers of nuclei within each tumor stage. **D)** Correlation between percentage of immune cells from DroNc-seq data and immune cell percentage from multiplex immunofluorescence (mIF) data for 13 samples. Samples are colored by tumor stage. Spearman correlation was used to determine the correlation coefficient *R* and *p*-value. **E)** Correlation between percentage of immune cells from DroNc-seq data and estimated immune score from bulk RNA-seq data for 44 samples. Samples are colored by tumor stage. Spearman correlation was used to determine the correlation coefficient *R* and *p*-value.

The strength of the snRNA-seq method is the possibility to analyze frozen tumors from patients with known clinical outcomes. Among the patients included in this study, 15 received at least five instillations with BCG. Although based on few samples and a low number of identified lymphocytes, we found that a lower fraction of lymphocytes in pre-BCG tumors was significantly associated with progression to MIBC (*p*=0.018). Additionally, no significant associations between specific cell populations and response to BCG or chemotherapy were observed.

Analyzing NMIBC and MIBC tumors separately did not reveal additional separation of tumors and clustering of non-epithelial cells across tumors was still driven by patient origin (*results not shown*). Attempts to correct for the sample-specific variation within non-epithelial cells yielded over-correlated, uninformative UMAPs and graph-based clusterings, possibly due to the low number of non-epithelial nuclei. Consequently, we did not investigate the non-epithelial cells further in this present work.

### Intra-tumor subtype heterogeneity accessed at single cell resolution

We focused our analysis on the epithelial (cancer cell) compartment for each tumor individually. Bulk RNA-seq data was available for 44 of the tumors and they were classified according to the six consensus classes of MIBC^2^ or the four UROMOL2021 classes of NMIBC^3^ depending on tumor stage. To assign a cancer cell phenotype to each epithelial cell and explore intra-tumor subtype heterogeneity at the single nucleus level, we used the bulk classification systems to classify all epithelial nuclei from tumors having >100 nuclei and at least 40% of the respective classifier genes expressed (NMIBC: 29,484 nuclei for 17 tumors; MIBC: 9,523 nuclei from 10 tumors). Tumors displayed varying levels of intra-tumor subtype heterogeneity with the dominating class at the single nucleus level constituting 36%-97% of all nuclei (mean 69%; **Figure 3A-B**), indicating that a discrete subtype assignment from bulk RNA-seq data may lack biological granularity. The dominating class at single nucleus level was consistent with the transcriptomic class from bulk RNA-seq in 79% (11/14) of NMIBC tumors (not considering class 2b tumors) and 67% (6/9) of MIBC tumors. We note that the low proportion of class 2b assignments at the single nucleus level for tumors classified as class 2b at the bulk RNA-seq level probably are due to the fact that class 2b tumors are characterized by a high immune infiltration of the tumor microenvironment which is not captured when looking at single nuclei from cancer cells only.

**Figure 3.**
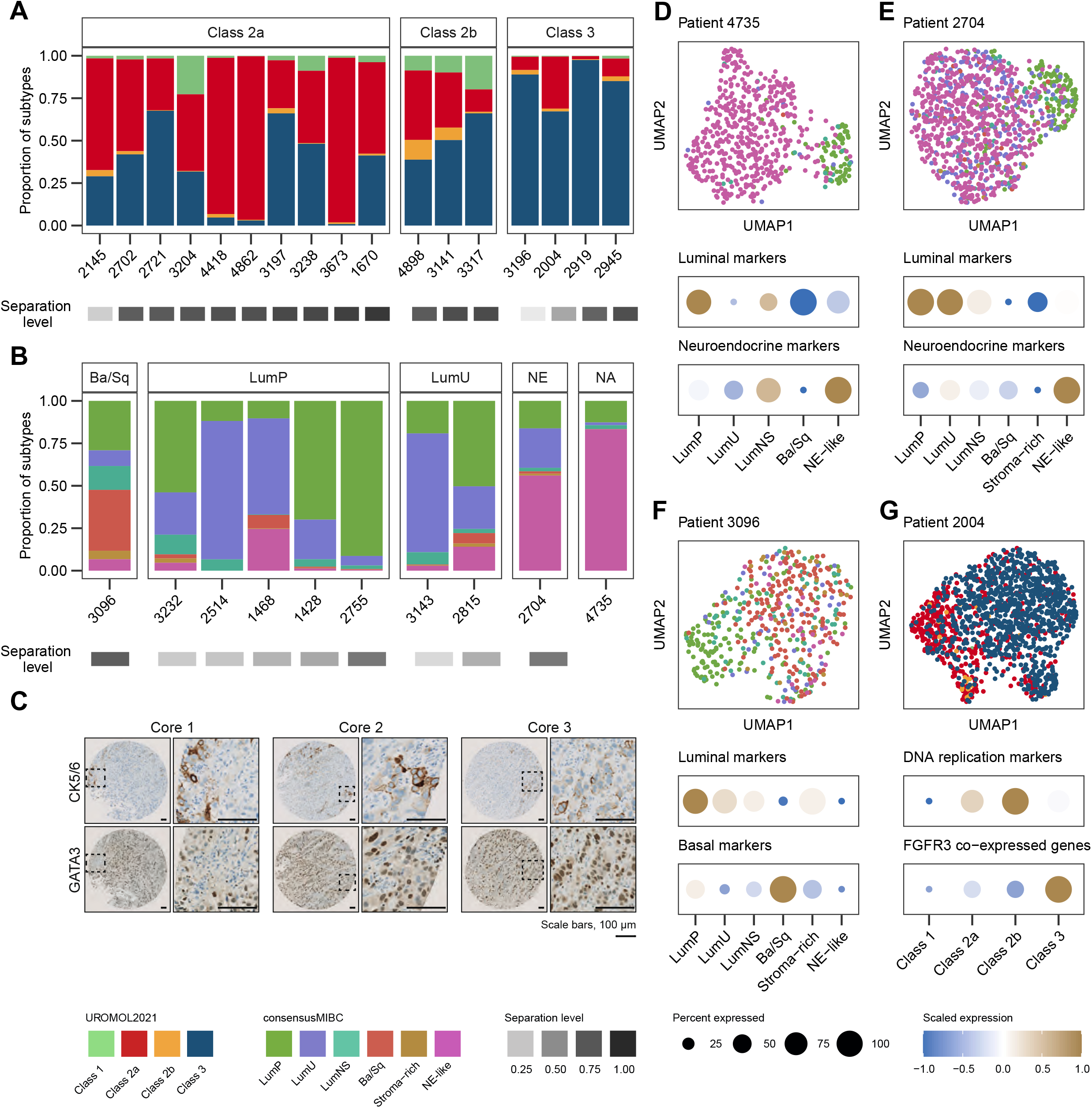
Intra-tumor subtype heterogeneity accessed at single cell resolution. **A)** Relative proportion of single nucleus subtype classifications of epithelial cells within NMIBC tumors separated by the subtype from bulk RNA-sequencing. Classification according to the UROMOL2021 system. Samples are sorted by increasing separation levels from the bulk classification. **B)** Relative proportion of single nucleus subtype classifications of epithelial cells within MIBC tumors separated by the subtype from bulk RNA-sequencing. Classification according to the consensusMIBC system. Samples are sorted by increasing separation levels from the bulk classification. The tumor from patient 4735 was not classified as no bulk RNA-sequencing was available. **C)** Immunohistochemical staining for CK5/6 and GATA3 of three areas (tissue microarray cores) from the tumor from patient 1468. **D-G)** UMAP visualization of selected tumors colored by single nucleus classifications (top) and dot plots showing the mean expression of subtype-specific gene signatures within each subtype population (bottom). Number of nuclei within each class for **D)** patient 4735; LumP: 71, LumU: 9, LumNS: 13, Ba/Sq: 1, NE-like: 465, **E)** patient 2704; LumP: 163, LumU: 234, LumNS: 21, Ba/Sq: 14, Stroma-rich: 11, NE-like: 564, **F)** patient 3096; LumP: 139, LumU: 44, LumNS: 67, Ba/Sq: 171, Stroma-rich: 24, NE-like: 32 and **G)** patient 2004; class 1: 7, class 2a: 435, class 2b: 24, class 3: 955.

We investigated the intra-tumor heterogeneity further using immunohistochemical (IHC) staining of the MIBC tumor from patient 1468. Based on the single nucleus analysis, this tumor was mainly luminal of origin but it also consisted of a minor fraction of nuclei classified as Ba/Sq (**Figure 3B**). Interestingly, IHC staining of three tissue microarray cores revealed areas with malignant cells positive for the basal marker cytokeratin 5/6 (CK5/6) while the majority of malignant cells in all cores were GATA3 positive, documenting the presence of both luminal and basal features simultaneously (**Figure 3C**).

Next, we performed graph-based clustering and UMAP visualization of the tumors individually and found cases where the transcriptomic classes were a major driver of the clustering (**Figure 3D-G**; **Supplementary Figure S1B-C**). A histological neuroendocrine tumor (patient 4735) consisted of two major populations: nuclei classified as neuroendocrine-like (NE-like) and nuclei classified as luminal papillary (LumP). This subtype-dependent division was consistent with the expression of marker genes for neuroendocrine tumors (**Figure 3D**). In agreement with this, a tumor classified as NE-like from the bulk RNA-seq (patient 2704) showed a major cluster of mixed nuclei classified as either NE-like or luminal unstable (LumU) and a minor cluster of LumP classification (**Figure 3E**). Tumors from both patient 3096 and patient 2004 consisted of a major cluster in accordance with the transcriptomic class from bulk RNA-seq (Ba/Sq and class 3, respectively), along with minor clusters of LumP and class 2a, respectively, further emphasizing the presence of intra-tumor subtype heterogeneity beyond discrete bulk classifications (**Figure 3F-G**).

Next, we applied SCENIC^17^, a method to reconstruct regulons (gene regulatory networks consisting of transcription factors and their target genes) and infer regulon activity scores for each nuclei using all nuclei with >500 genes and tumors with at least 50 nuclei (21,583 nuclei from 39 tumors). We explored regulon activities associated with the single nucleus subtype classifications for NMIBC and MIBC tumors to further support the single nucleus subtype classifications. In agreement with findings from the UROMOL2021 study^3^, class 3 nuclei showed a significantly higher activity of the TP63, KDM5B and RARG regulons compared to the other UROMOL2021 classes (**Supplementary Figure S2A**). For MIBC, luminal tumors showed higher activity of e.g. the GATA3 and FOXA1 regulons and stroma-rich tumors showed a higher activity of the PGR and STAT3 regulons, as previously observed^2^ (**Supplementary Figure S2B**). Furthermore, subtype-specific regulon activities were also associated with single nucleus subtype classifications when analyzing selected tumors individually (**Figure 4A-D**).

**Figure 4.**
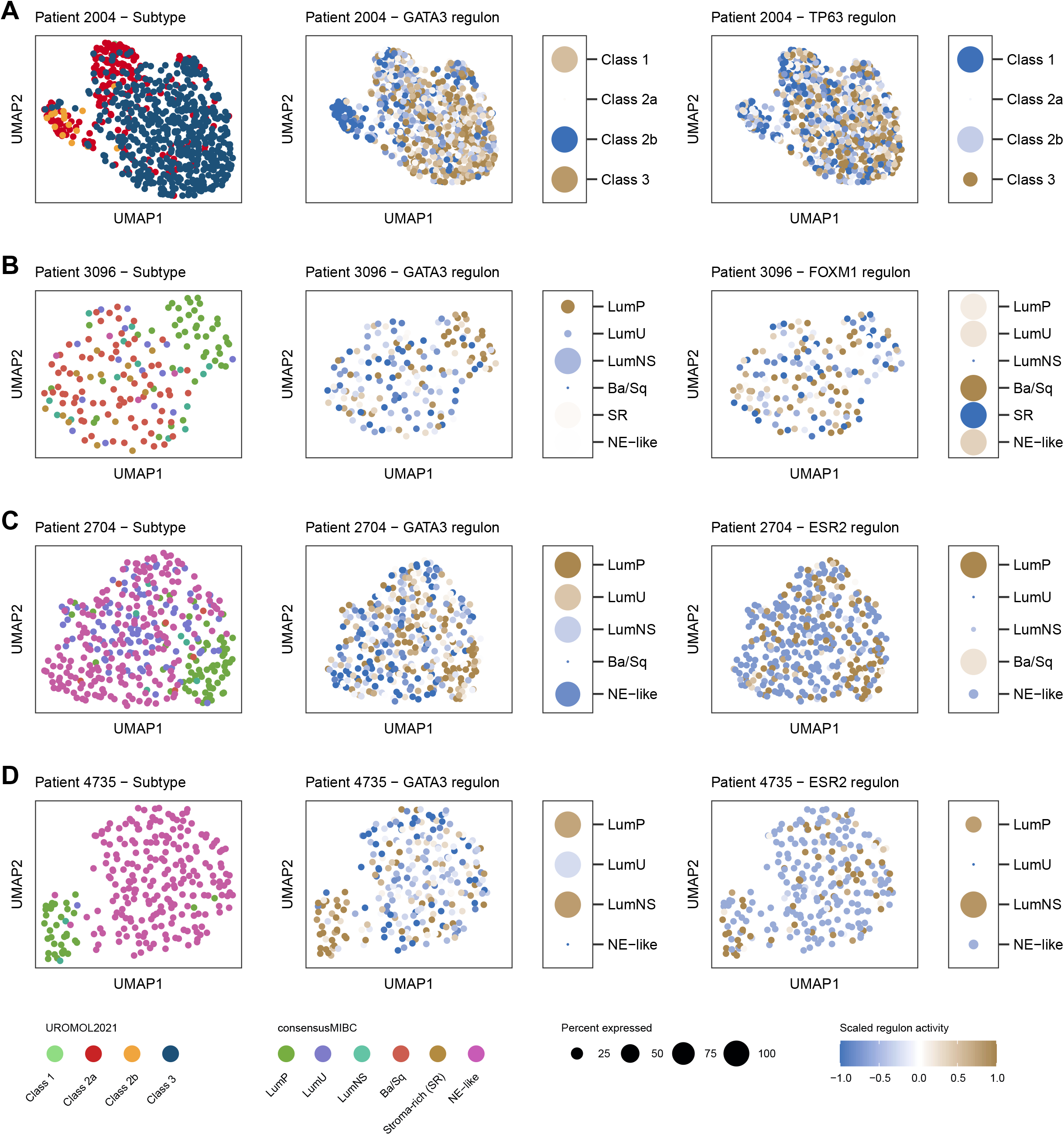
Regulon activity scores of single nuclei. **A-D)** UMAP visualization of selected tumors colored by single nucleus subtype classifications (left) and activity scores for selected subtype-specific regulons (middle, right). Dot plots of mean regulon activities within each subtype population are shown to the right of each regulon-colored UMAP. Number of nuclei within each class for **A)** patient 2004; class 1: 2, class 2a: 202, class 2b: 16, class 3: 515, **B)** patient 3096; LumP: 45, LumU: 15, LumNS: 14, Ba/Sq: 72, Stroma-rich: 14, NE-like: 4, **C)** patient 2704; LumP: 66, LumU: 88, LumNS: 6, Ba/Sq: 6, NE-like: 205 and **D)** patient 4735; LumP: 35, LumU: 1, LumNS: 2, NE-like: 220.

### Intra-tumor subtype heterogeneity assessed by spatial transcriptomics

To explore potential spatial organization of intra-tumor subtype heterogeneity, we performed spatially resolved whole transcriptomics of four bladder samples including two T1 tumors (patient 2919 and 3197) and two T2-4 tumors (patient 4735 and 3143) using the 10x Visium Spatial platform. We classified each spot using the appropriate bulk classification system, knowing that the assigned classifications constitute an average of the cells present in the different spots (**Figure 5A-D**). The tumors from patient 2919 and 4735 showed high subtype homogeneity at the single nucleus level and this was confirmed at the spatial transcriptomics level (**Figure 5A** and **5C**). Although being class 2a at the bulk level, the tumor from patient 3197 showed a high proportion of low-risk class 3 at the single nucleus level and low-risk classifications (class 1 and 3) were also dominant with spatial transcriptomics (**Figure 5B**). Finally, the spots for patient 3143, which was classified as LumU at the bulk level, were mainly classified as LumNS. This could be explained by tumor heterogeneity as only a smaller tissue section was analyzed using spatial transcriptomics (**Figure 5D**). However, a minor fraction of LumNS classification was observed at the single nucleus level as well (**Figure 3B**). Furthermore, we utilized the anchor-based integration workflow in Seurat to deconvolute the cellular composition of each spot using the same snRNA-seq reference of human bladder tumors^12^ as previously. In general, we observed that areas with higher scores for non-epithelial cell compartments were classified as class 2b for NMIBCs or stroma-rich for MIBCs (**Figure 5A-D**).

**Figure 5.**
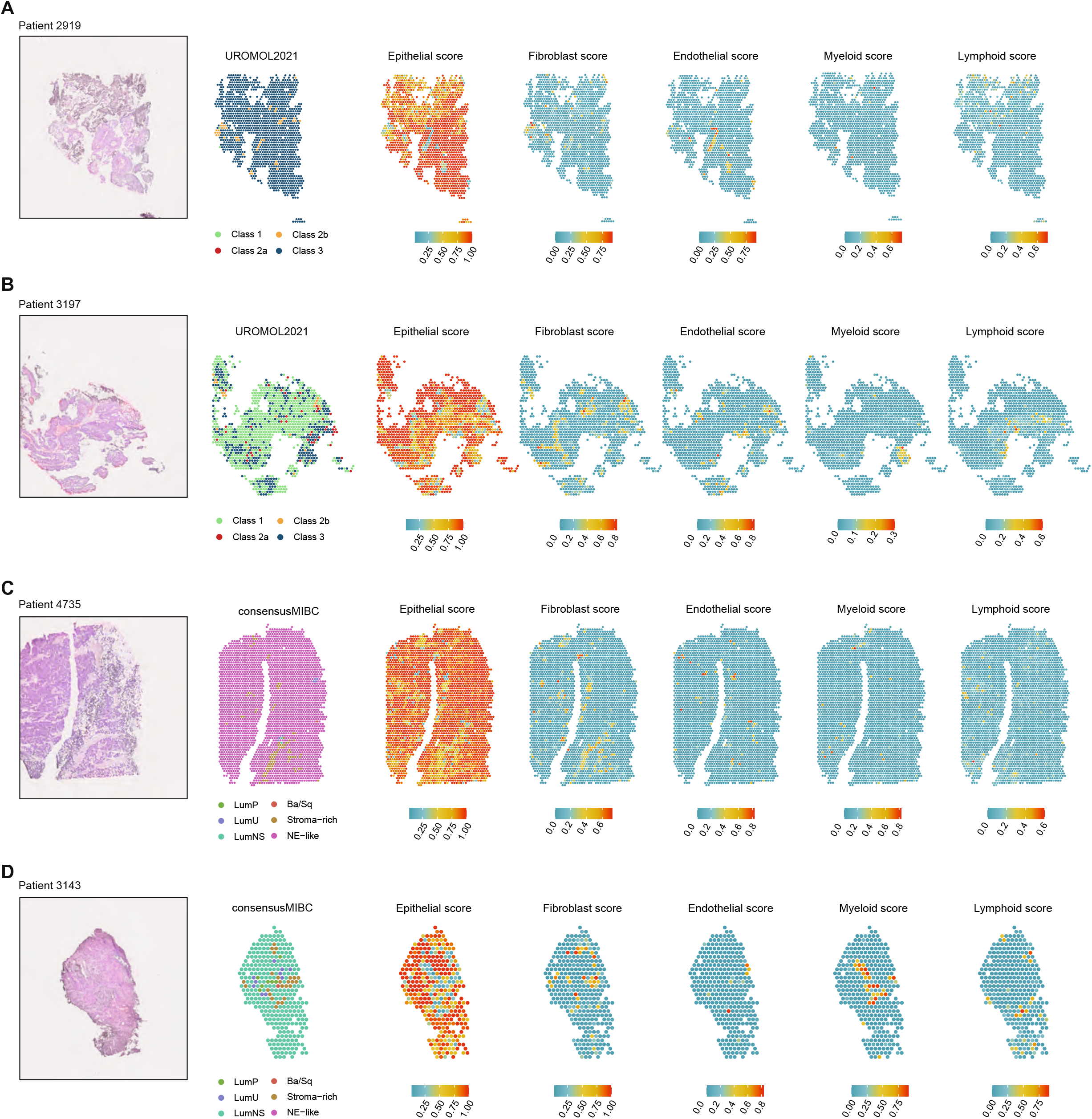
Intra-tumor subtype heterogeneity assessed by spatial transcriptomics. **A-D)** Images of H&E staining of tissue areas (left) from the four tumors analyzed using 10x Visium Spatial. Spots are colored by subtype classifications (middle) and cellular composition scores (right).

### Delineating intra-tumor subtype heterogeneity from bulk tumor samples

Based on this knowledge of intra-tumor subtype heterogeneity at the single nucleus level, we next investigated the impact of the presence of high-risk subtypes in tumor subclones that may not be identified from bulk subtype assignments. To approximate intra-tumor subtype heterogeneity from bulk transcriptomic profiles, we used the deconvolution tool Weighted In-Silico Pathology (WISP)^18^ on the UROMOL2021 (n=505) and TCGA cohorts (n=406). WISP calculates pure population centroid profiles and provides a weighting of each sample between the transcriptomic subtypes. As observed at the single nucleus level, tumors displayed a varying level of intra-tumor subtype heterogeneity on the bulk tumor level as well (**Figure 6A-B**). WISP weights correlated with the proportion of class assignments at the single nucleus level, especially for class 2a (**Figure 6C**; **Supplementary Figure S3**). Finally, we explored the clinical impact of the level of class 2a assignments, as class 2a is the molecular high-risk subtype of NMIBC tumors. When only considering tumors with a bulk class 2a assignment, we found that a high class 2a weight (estimated from the bulk RNA-seq data using WISP) was significantly associated with worse progression-free survival (PFS; *p*=0.0005; **Figure 6D**). Notably, a high class 2a weight was still associated with worse PFS when adjusted for the EAU clinical risk assessment (HR=2.97, 95%CI=1.22-7.23, *p*=0.017) and EORTC risk score (HR=2.58, 95%CI=1.06-6.29, *p*=0.036; **Supplementary Table S2**).

**Figure 6.**
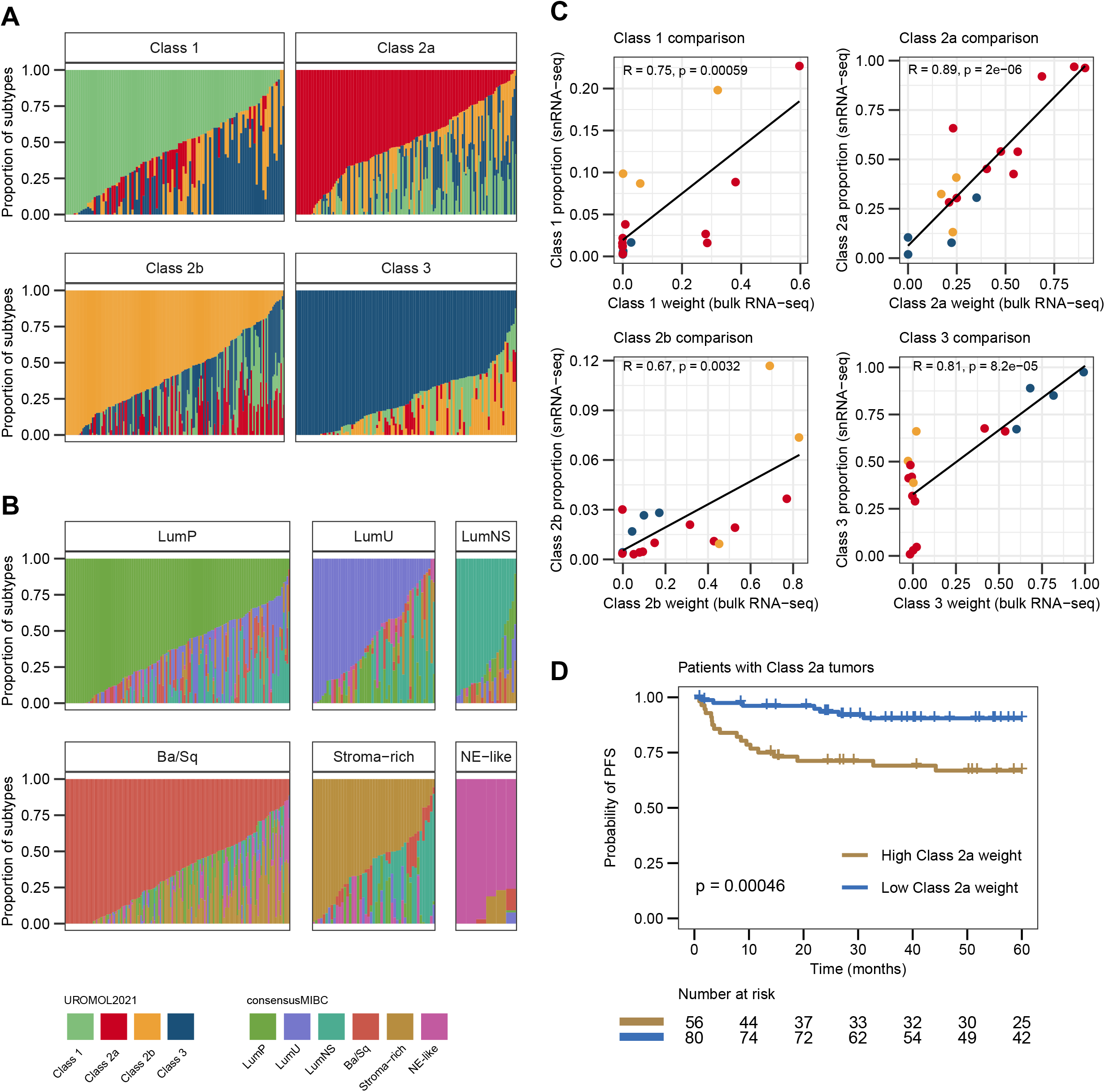
Delineating intra-tumor subtype heterogeneity from bulk tumor samples. **A)** WISP weights of the four UROMOL2021 classes for 505 tumors from the UROMOL2021 cohort separated by bulk subtype assignment. **B)** WISP weights of the six consensus classes of MIBC for 406 tumors from the TCGA cohort separated by bulk subtype assignment. **C)** Correlation between bulk WISP weights and DroNc-seq class proportions for each of the UROMOL2021 classes. Samples are colored by bulk subtype assignment. Pearson correlation was used to determine the correlation coefficient *R* and *p*-value. **D)** Kaplan-Meier plot of progression-free survival (PFS) for 136 patients with class 2a tumors stratified by class 2a weight from WISP analysis (log-rank test). An optimal cutpoint for the class 2a WISP weight according to time to progression was determined using the R package survminer (cutpoint=0.5562).

## Discussion

Here we performed snRNA-seq analysis of 48 tumors from patients across non-muscle invasive and muscle invasive disease and four tumors were additionally analyzed using spatial transcriptomics. By analyzing frozen biobanked samples, we were able to compare our results to previously generated total RNA-sequencing and mIF/IHC staining for 44 and 13 of the tumors, respectively. We demonstrated that tumors from both early and more advanced disease stages display large intra-tumor subtype heterogeneity and that the level of subtype heterogeneity could be estimated from both single nucleus and bulk RNA-seq data with a high concordance between the two. Notably, we showed that a high class 2a weight estimated from bulk RNA-seq data using WISP was associated with worse outcome in patients with molecular high-risk class 2a tumors, indicating that discrete subtype assignments from bulk RNA-seq data may lack biological granularity - i.e. high-risk subclones will be overlooked in bulk analysis. In line with our observations, a previous study of scRNA-seq of three murine and two human MIBC bladder tumors provided initial evidence of intra-tumor subtype heterogeneity as tumors consisted of multiple transcriptomic signatures derived from subpopulations of cells of different lineages^9^. This supports the potential use of continuous classification scores instead of discrete class assignments for improved risk stratification of patients. However, the use of continuous scores to separate patients need to be tested and validated in future studies to explore clinical utility further.

In this current study, we applied the consensusMIBC and UROMOL2021 bulk classification systems to snRNA-seq data without any optimization. Optimally, bulk classification systems should be adjusted when applied to sparse single cell data. A previous study on breast cancer developed a compatible method for intrinsic molecular subtyping according to the PAM50 gene signatures for scRNA-seq and used the scRNA-seq data to refine the molecular subtypes of breast cancer^19^. Furthermore, another recent study utilized findings from scRNA-seq of 63 colorectal tumors to refine the bulk consensus molecular subtype (CMS) classification of colorectal cancer^20^. Both studies highlight how single cell data can improve and increase the biological granularity of subtyping. However, due to the sparsity of the DroNc-seq data presented here, we chose not to develop new classification systems compatible for snRNA-seq data and did not refine the bulk classification systems of BC further. Observed inconsistency between the dominating class at single nucleus level and transcriptomic class from bulk RNA-seq may therefore be explained by several levels of heterogeneity (different parts of the tumor tissue goes to the various analyses) and methodological differences, including the technical challenges of applying a bulk classification system to single nucleus data. However, we did find an overall good concordance between bulk and single nucleus classifications. Furthermore, analyzing tumors individually revealed cases where the transcriptomic classes were a major driver of the clustering. Some of the most prominent cases included a histological neuroendocrine tumor (patient 4735) and a tumor classified as NE-like at the bulk RNA-seq level (patient 2704). This indicates that the sparse DroNc-seq data more easily captures the large biological difference between the NE-like and luminal subtypes compared to smaller differences within more pure luminal tumors.

The major advantage of using snRNA-seq over scRNA-seq is the ability to analyze frozen tumors where clinical outcomes of the patients are known. Although we did not perform a thorough exploration of the non-epithelial tumor compartments as it constituted a minor fraction of the analyzed nuclei, we found that a lower fraction of lymphocytes in pre-BCG tumors from 15 patients treated with a minimum of five BCG instillations was significantly associated with progression to MIBC. This is in concordance with previous findings based on mIF staining where high infiltration of cytotoxic T cells was associated with a lower risk of progression^15^. Furthermore, a previous study showed that the level of infiltration and extent of T cell exhaustion were associated with outcome and response to BCG treatment^16^. We observed that the highly infiltrated subtypes, i.e. class 2b for NMIBCs and stroma-rich for MIBCs, were not captured when looking at single nuclei from cancer cells only, as these subtypes are mainly characterized by a highly infiltrated tumor microenvironment. Therefore, given the clinical impact of infiltration, the level of infiltration should still be considered if bulk classifications systems are refined by epithelial cell phenotypes observed in snRNA-seq or scRNA-seq data.

Finally, our comparison of methods showed a better performance of the 10x Chromium platform with a notable increase in data per nuclei compared to the DroNc-seq system. This could be due to higher capture rate, better error suppression in nucleus barcodes or the utilization of elastic beads when using 10x Chromium. However, while the 10x Chromium platform is less time-consuming and a more standardized system, the DroNc-seq system has lower running costs and was therefore selected for this exploratory study to increase the cohort size.

In conclusion, we demonstrated that intra-tumor subtype heterogeneity is an important biological feature of human bladder tumors that may have an impact on clinical outcome. Continuous subtype scores may have the potential to further refine the biological characterization of tumors and clinical stratification of patients and should be considered when designing clinical trials investigating the clinical impact of molecular subtyping in BC.

## Acknowledgements

We would like to thank all technical personnel at the Department of Molecular Medicine, Urology and Oncology, Aarhus University Hospital, for sample handling and processing.

## Author contributions

Conceptualization, S.V.L., S.S.S., I.N., L.D.; Methodology, S.S.S., I.N., L.D.; Formal Analysis, S.V.L., S.S.S., P.L.; Investigation, S.V.L., S.S.S., I.N., P.L., M.K., F.P., T.S., L.D.; Resources, J.B.J., L.D.; Data Curation, S.V.L., S.S.S., P.L., M.K.; Writing - Original Draft, S.V.L., S.S.S., L.D.; Writing - Review and Editing, all authors; Visualization, S.V.L.; Supervision, L.D.; Project Administration, L.D.; Funding Acquisition, L.D.

## Competing interests

Lars Dyrskjøt has sponsored research agreements with C2i, AstraZeneca, Natera, Photocure, and Ferring; has an advisory/consulting role at Ferring and UroGen; and is Chairman of the Board in BioXpedia A/S.

Jørgen Bjerggaard Jensen is proctor for Intuitive Surgery; is a member of advisory board for Olympus Europe, Ambu, Cepheid, and Ferring; and has sponsored research agreements with Medac, Photocure ASA, Cepheid, and Ferring.

## Data availability

The raw sequencing data generated in this study are not publicly available as this compromise patient consent and ethics regulations in Denmark. Processed non-sensitive data are available upon reasonable request from the corresponding author.

## Funding information

Independent Research Fund Denmark, The Novo Nordisk Foundation, Aarhus University (AUFF NOVA), The Leo & Anne Albert Institute for Bladder Cancer Care and Research.

---

**Supplementary Figure S1. Non-epithelial cell proportions and tumor-specific UMAPs. A)** Relative proportions of non-epithelial cell types within three samples analyzed on two platforms; 10x Chromium (2004, n=161; 2514, n=92; 4735, n=91) and DroNc-seq (2004, n=104; 2514, n=155; 4735, n=25). **B)** UMAP visualization of NMIBC tumors colored by single nucleus classifications according to the UROMOL2021 system. **C)** UMAP visualization of MIBC tumors colored by single nucleus classifications according to the consensusMIBC system.

**Supplementary Figure S2. Regulon activities within nuclei divided by subtype classifications. A)** Scaled regulon activities in each subtype population of NMIBC tumors. Top 15 regulons with an adjusted *p*-value<0.01 within each subtype are shown. Regulons included in the UROMOL2021 study are shown in bold font. **B)** Scaled regulon activities in each subtype population of MIBC tumors. Top 10 regulons with an adjusted *p*-value<0.01 within each subtype are shown. Regulons included in the consensusMIBC study are shown in bold font.

**Supplementary Figure S3. Comparison between single nucleus and bulk classifications for MIBC tumors. A)** Correlation between bulk WISP weights and DroNc-seq class proportions for each of the consensusMIBC classes. Samples are colored by bulk subtype assignment. Pearson correlation was used to determine the correlation coefficient *R* and *p*-value.

**Supplementary Table S1**: Clinical and molecular information of analyzed samples.

**Supplementary Table S2**: Univariate- and multivariable cox regression analysis.

## Methods

### Tumor samples

A total of 48 bladder tumor samples (10 Ta, 13 T1, 25 T2-4) were obtained from transurethral resection of the bladder (TURB) or radical cystectomy (**Supplementary Table S1**). One sample was processed directly from surgery and 47 samples were embedded in O.C.T., frozen in liquid nitrogen and stored at −80 °C. All patients provided written informed consent to participate in future research projects and the study was approved by the National Committee on Health Research Ethics (#1706291 and #1708266).

### Nuclei isolation and snRNA-seq using an optimized DroNc-seq protocol

We performed snRNA-seq of bladder tumors using an optimized DroNc-seq protocol previously described^13^. Briefly, frozen tumors were sectioned and the fresh tumor was dissected using pestles and incubated in IgePal lysis buffer^13^ to isolate single nuclei. Following breakage of the cytoplasmic membrane the nuclei were isolated, washed and evaluated under a fluorescence microscope using DAPI staining. The nuclei were diluted to a final concentration of 453,000 nuclei per ml and subjected to the Dolomite Bio platform for droplet generation. Droplet breakage, reverse transcription (RT) and cDNA amplification was performed as described by Habib and colleagues^21^. Tagmentation and amplification was performed using Nextera XT DNA Library Preparation Kit (Illumina, cat #FC-131-1096) and 600 pg input of each sample. SPRI cleanup and quality assessment of cDNA and final DNA libraries was performed^13^ and single nuclei libraries were sequenced on the Illumina NovaSeq 6000 platform^13^.

### snRNA-seq data processing and quality control

Processing of the FASTQ files was done as described in the Drop-Seq Core computational Protocol with a few supplementary steps. First, we sorted the reads and kept only the ones where the barcodes were correctly formed, i.e. the molecular barcode followed by a V (i.e. a A, C or G base) and then the polyT. Then we ran the pipeline as described. To summarize: tag the biological read (read2) with the barcode and the unique molecular identifier (UMI), filter reads with low quality barcodes, trim 5’ primer sequence, trim 3’ polyA, align the biological read with STAR, add gene/exon and other annotation tags and finally repair substitution errors or indel errors.

For each experiment, we then calculated the expression per barcode for all barcodes with a minimum of 150 genes expressed. Next, we generated a whitelist of barcodes for each tumor by selecting the *x* number of barcodes with the highest number of genes expressed, where *x* was based on the theoretical, expected number of nuclei from the lab procedure (**Supplementary Table S1**). We then created different tags where non-white-listed barcodes were changed to a white-listed barcode if the hamming distance between the two barcodes was 0 (H0), less or equal to 1 (H1) or less or equal to 2 (H2). Finally, for each tag, we recalculated the expression of each gene for all the barcodes in the white-list.

We used the R package Seurat^22^ v3.2.0 to further process and analyze the snRNA-seq data. The preprocessed count matrices for all samples were initially merged into a combined count matrix (51,604 genes x 117,653 nuclei). We estimated the level of ambient RNA contamination using the R package DIEM^23^ v2.3.0 with default parameters. Nuclei with <200 genes expressed were fixed as “debris” (n=9,105) and nuclei with an estimated debris score >0.5 were removed (n=14,382). Next, we removed nuclei with >10% of UMIs mapping to mitochondrial genes (n=19,545) and predicted potential doublets using the R package DoubletFinder^24^ v2.0.3 (n=3,729). Each sample was processed individually as required, and DoubletFinder was run with default settings and an expected doublet rate of 5%. Finally, the top 5% and bottom 10% of nuclei based on the number of expressed genes (<267 and >1,044 genes) and number of UMIs (<326 and >1,536 UMIs) were removed (n=11,840). This resulted in 59,052 nuclei, which were used for downstream analysis (1,230 nuclei per sample on average).

### snRNA-seq using 10x Chromium

Three samples were additionally analyzed using the 10x Chromium platform. Single nuclei suspensions were loaded onto the Chromium Next GEM Chip G for droplet generation followed by library preparation according to the manufacturer’s recommendations (Chromium Next GEM Single Cell 3ʹ (PN-1000128), 10x Genomics). In short, single nuclei were encapsulated with Gel Beads-in-Emulsion in the 10x Chromium controller followed by nuclei lysis and RT creating barcoded, full-length cDNA. Then, amplification, fragmentation, end repair, A-tailing, adaptor ligation and sample indexing were performed. An input of 5,000 single nuclei were loaded to each channel with a targeted nuclei recovery rate of 3,000 nuclei. Libraries were sequenced on an Illumina NovaSeq 6000 platform.

Data was processed with Cell Ranger v7.0.0 (10x Genomics) and mapped using a pre-mrna reference (GRCh38-2020-A) including introns. In total, we obtained data from 7,214 nuclei. Using the R package Seurat^22^ v3.2.0, we removed nuclei with >10% of UMIs mapping to mitochondrial genes (n=1,022), potential doublets as described above (n=309) and nuclei with <200 or >6000 genes (n=125). Thereby, 5,758 nuclei remained for downstream analysis.

### Cell type annotation

The R package SingleR v1.0.6 was used for cell type annotation^25^. We removed genes expressed in <10 nuclei and obtained log-normalized expression values using NormalizeData from Seurat. As reference dataset, we chose the recently published dataset from Gouin et al. containing snRNA-seq of 25 treatment-naive muscle invasive bladder tumors (GSE169379)^12^. All nuclei in the downloaded dataset were pre-labelled as either epithelial, fibroblast, endothelial, lymphoid or myeloid, and we log-normalized the data using NormalizeData from Seurat. As recommended when using a single cell reference dataset, SingleR was run in the Wilcoxon rank sum test marker detection mode (de.method=”wilcox”) and with 10 marker genes from each pairwise comparison between labels (de.n=10). We removed nuclei with a “delta median value” (the difference between the score for the assigned label and the median across all labels for that nuclei) below 0.05 to filter assignments of low quality (n=10,485). The three samples analyzed using 10x Chromium were annotated using the same approach as described above. In total, 5010 nuclei remained after removing low-quality assignments with a delta median value below 0.05.

### Transcriptomic classification of single nuclei

All epithelial nuclei were classified according to the four UROMOL2021 classes^3^ or six consensus classes of MIBC^2^. The transcriptomic classification of nuclei was performed for each tumor individually: Genes expressed in <10 nuclei were removed and the count data was log-normalized using NormalizeData in Seurat. Only tumors having >100 epithelial nuclei and at least 40% of the respective classifier genes expressed were included.

DNA replication markers and FGFR3 co-expressed genes were extracted from Lindskrog et al.^3^, luminal- and basal markers were extracted from Kamoun et al.^2^ and neuroendocrine markers were extracted from Batista da Costa et al.^26^. For all gene signatures, we calculated the mean expression of the marker genes for each nuclei.

### Dimensionality reduction, clustering and visualization

Using the merged and filtered count matrix, we kept genes with non-zero expression in >=10 nuclei and performed log-normalization using NormalizeData in Seurat (default settings). The 2,000 most variable genes were selected, and the number of counts and the percentage of mapping to mitochondrial genes were regressed out using ScaleData. We performed dimensionality reduction of the variable genes by principal component analysis and selected the top 20 principal components as input for graph-based clustering (resolution=0.5) and UMAP visualization.

We chose not to correct for sample-specific batch effects as 1) batch correction algorithms yielded over-correlated, uninformative UMAP clusterings and 2) we believe that epithelial tumor cells have genuine sample-specific variation as previously shown^27^, and the epithelial compartment constituted the bulk of the tumors.

For the analysis of individual samples, we applied the workflow as for the merged dataset, except that we used the top 10 principal components as input for graph-based clustering (resolution=0.5) and UMAP visualization.

### SCENIC regulon analysis

We applied the SCENIC workflow v0.11.2 (pySCENIC) to reconstruct regulons (transcription factors and their target genes) and estimate regulon activity within single nuclei. Using the merged dataset of 48,567 annotated nuclei, we removed genes expressed in <1% of nuclei and nuclei with <500 genes expressed. Furthermore, we excluded tumors with <50 nuclei, resulting in a dataset of 9,852 genes and 21,583 nuclei from 39 tumors which was saved as a loom file. The SCENIC analysis was performed with a curated list of human transcription factors^28^ and a cisTarget database of motif enrichments 10 kilobases up and downstream of transcription start sites (motif database version: hg38 mc9nr, motif annotations: v9) as described in Van de Sande et al.^17^. AUCell scores were calculated for each of the inferred regulons. The FindAllMarkers function within Seurat was used to identify regulons with different activity between subtypes (settings: test.use=”wilcox”, only.pos=T, logfc.threshold=0.005).

### Bulk RNA-sequencing data

Bulk RNA-sequencing (RNA-seq) data was available for 44 tumors. For 16 of the tumors, library preparation was performed using ScriptSeq (EpiCentre) followed by sequencing on Illumina HiSeq 2000 or KAPA RNA HyperPrep Kit (RiboErase HMR; Roche) followed by sequencing on Illumina NovaSeq 6000^16^. For the remaining 28 tumors, RNA-seq was performed using the QuantSeq kit FWD HT kit (Lexogen) followed by sequencing on Illumina NextSeq 500^14^. NMIBC and MIBC tumors were classified according to the four UROMOL2021 transcriptomic classes^3^ and the six consensus subtypes of MIBC^2^, respectively. We estimated immune cell populations from the bulk RNA-seq data using established gene expression signatures^29^ (for CD4 T cells^30^) as in Rosenthal et al^31^. A score for each immune cell population was calculated as the mean expression of all marker genes for the given cell type and a total immune score was defined as the mean of all individual immune cell population scores.

### Weighted In-Silico Pathology (WISP) analysis

We used the R package WISP v2.3 to deconvolute intra-tumor heterogeneity and class stability from bulk transcriptomic profiles^18^. Normalized expression data of 406 muscle invasive TCGA tumors were obtained using the R package TCGAbiolinks v2.22.4. Clinical information of the samples were obtained from Robertson et al^32^. Data was log2 transformed and all samples were classified according to the six consensus classes of MIBC^2^. Normalized expression data, clinical information and class assignments of 535 non-muscle invasive UROMOL tumors were obtained from Lindskrog et al^3^. Only tumors with a positive silhouette score were used for the WISP analysis (n=505). Pure population centroid profiles were calculated for both cohorts separately using the respective classification systems: 149 and 167 samples were kept as “pure” for the UROMOL2021 and TCGA cohort, respectively, and 50 top marker genes for each subtype were included in the centroid profiles. Using the centroid profiles, WISP weights were then calculated for each sample in the two cohorts and for the bulk transcriptomic profiles of the samples included in the current snRNA-seq study. Pearson correlation was used to evaluate association between WISP weights and class proportions from snRNA-seq data. For the survival analysis, an optimal cutpoint for the class 2a WISP weight according to time to progression was determined using the R package survminer v0.4.9 (cutpoint=0.5562).

### Immunofluorescence and immunohistochemical staining

Tissue microarrays were previously generated. Standard bright-field immunohistochemistry was used for the detection of GATA3 and CK5/6 and a tyramide signal amplification strategy was used for mIF detection of two antibody panels (panel 1: CD3, CD8, FOXP3; panel 2: CD20, CD68, CD163, HLA-A, HLA-B, HLA-C) as described in Taber al.^15^.

### Spatial transcriptomics

Four samples were analyzed using the 10x Visium Spatial platform. Sections of tissue were cut for methanol fixation, H&E staining and imaging followed by visium spatial gene expression library construction as described by the manufacturer (CG000160 and CG000239, 10x Genomics). In short, sections of tissue were methanol-fixated, H&E stained and imaged. Then, the tissue sections were permeabilized, RT were performed and cDNA were amplified and used for downstream library construction including fragmentation, end repair and A-tailing, adaptor ligation, sample indexing and quality assessment. Libraries were sequenced on an Illumina NovaSeq 6000.

Data was mapped to the reference genome GRCh38 using Space Ranger v1.2.0 (10x Genomics). Count matrices and images were loaded into R using Seurat v4.2.0 and STutility^33^ v1.1.1 for subsequent data filtering, normalization and visualization. We removed spatial spots with <500 genes expressed or >25% of reads mapping to mitochondrial genes. Furthermore, only genes detected in more than 5 spots were kept for further analysis. For each of the four samples, data normalization was performed using the variance stabilized transformation method implemented in the SCTransform function in Seurat. Spots were classified according to the four UROMOL2021 classes^3^ or six consensus classes of MIBC^2^.

### Statistical analysis

Survival analysis was performed using the Kaplan–Meier method and two-sided log-rank tests were used to compare survival curves (R packages survival v3.2.13 and survminer v0.4.9). All statistical and bioinformatic analyses were performed with R v3.6.1, except the SCENIC analysis which was performed with python v3.10.5.

