## Supplementary Figures and Tables for "Single nucleus and spatially resolved intra-tumor subtype heterogeneity in bladder cancer"

### Supplementary Figure S1

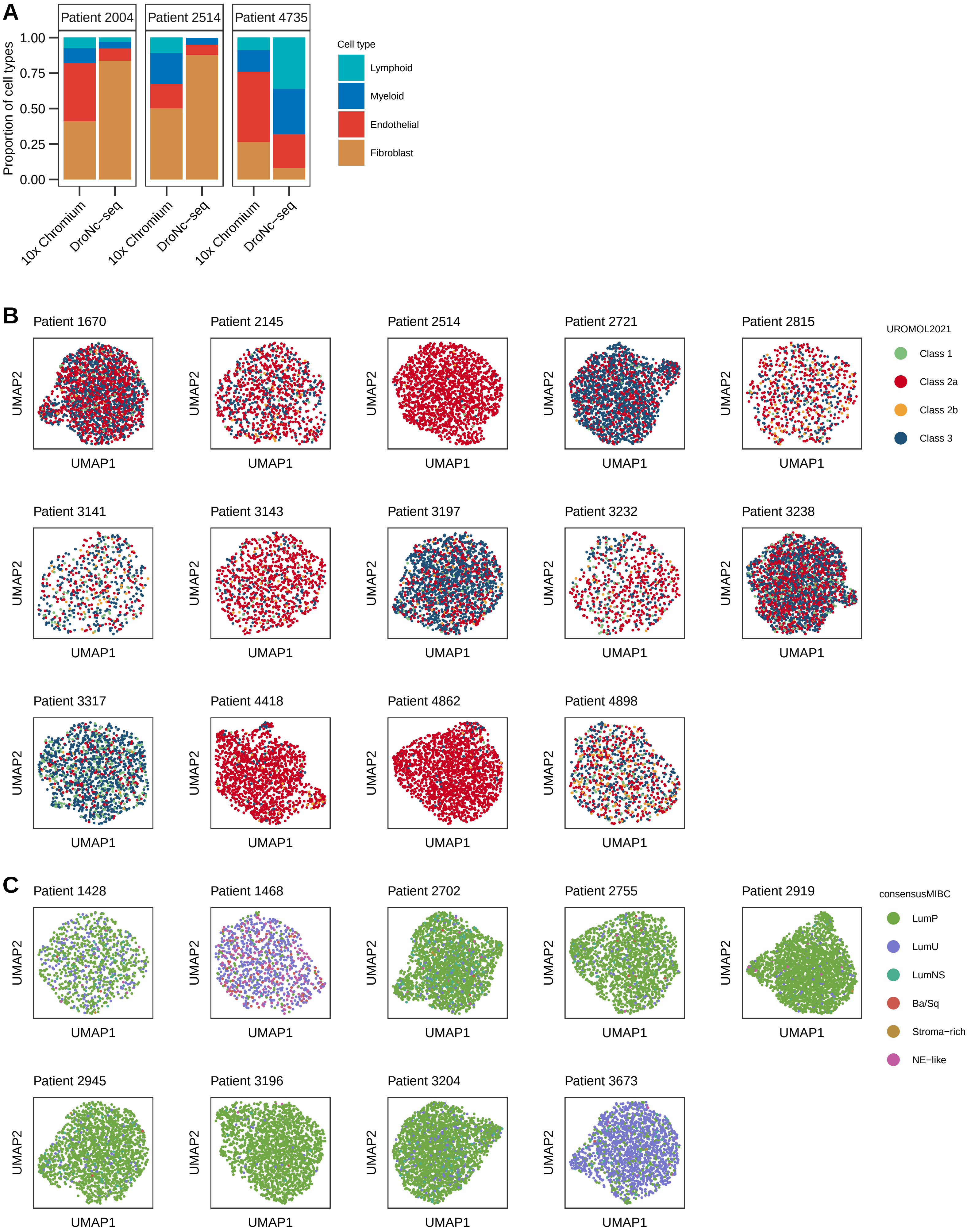

### Supplementary Figure S2

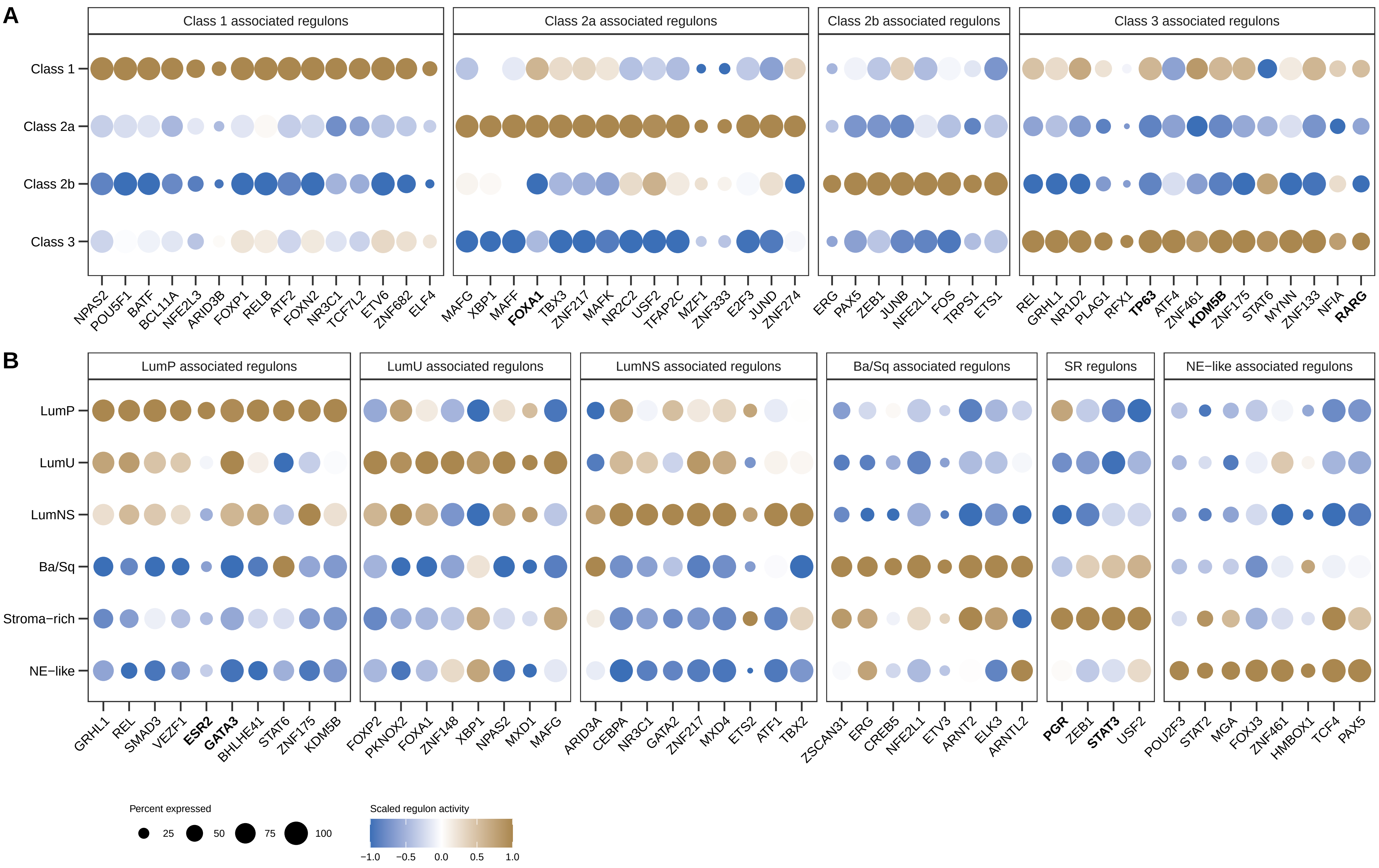

### Supplementary Figure S3

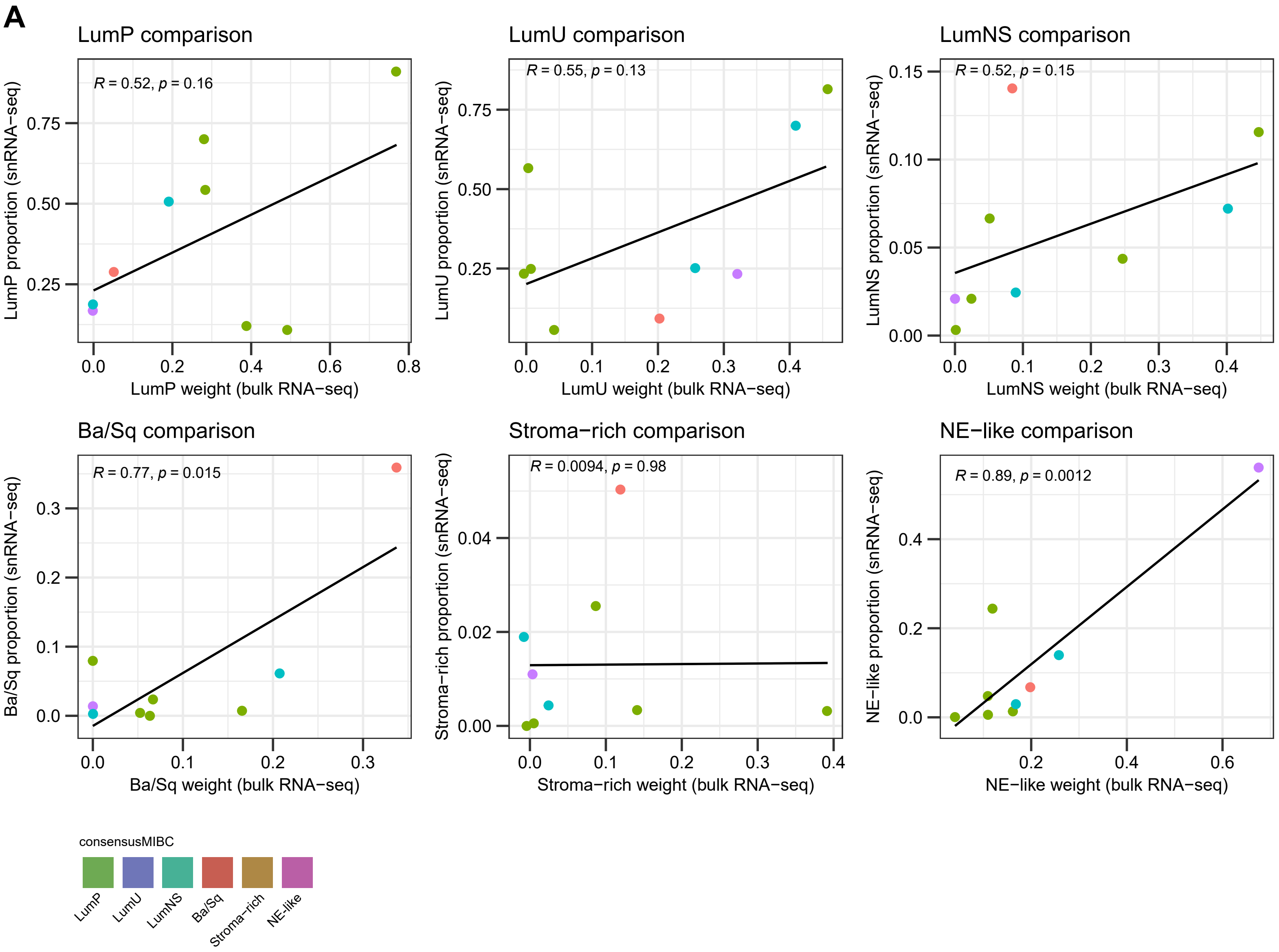

Supplementary Tabel S1: Clinical and molecular information of analyzed samples.

| PatientID | Gender | Tumor stage | Tumor grade | Bulk consensusMIBC class | Bulk UROMOL2021 class | Theoretical number of nuclei | Million uniquely mapped reads | #Nuclei after barcode processing | #Nuclei after QC filtering |
|---|---|---|---|---|---|---|---|---|---|
| 2945 | Male | Ta | Low grade | | Class 3 | 3,000 | 438 | 3,000 | 1,852 |
| 2721 | Male | Ta | Low grade | | Class 2a | 3,000 | 388 | 3,000 | 1,968 |
| 1841 | Male | Ta | Low grade | | Class 2a | 2,000 | 317 | 2,000 | 828 |
| 3779 | Male | Ta | High grade | | Class 2a | 3,000 | 283 | 3,000 | 186 |
| 4898 | Male | Ta | High grade | | Class 2b | 3,000 | 234 | 3,000 | 1,230 |
| 3238 | Male | Ta | High grade | | Class 2a | 3,000 | 202 | 3,000 | 2,424 |
| 3204 | Male | Ta | Low grade | | Class 2a | 3,000 | 193 | 3,000 | 2,606 |
| 2939 | Male | Ta | Low grade | | Class 3 | 3,000 | 113 | 3,000 | 995 |
| 3317 | Male | Ta | Low grade | | Class 2b | 3,000 | 72 | 3,000 | 1,473 |
| 1622 | Male | Ta | Low grade | | Class 2a | 1,000 | 29 | 792 | 44 |
| 2145 | Male | T1 | High grade | | Class 2a | 2,000 | 549 | 2,000 | 1,066 |
| 4059 | Male | T1 | High grade | | | 1,000 | 341 | 1,000 | 293 |
| 4862 | Male | T1 | High grade | | Class 2a | 4,000 | 340 | 4,000 | 1,805 |
| 3141 | Male | T1 | High grade | | Class 2b | 3,000 | 302 | 3,000 | 791 |
| 3196 | Male | T1 | High grade | | Class 3 | 3,000 | 292 | 3,000 | 1,989 |
| 3197 | Male | T1 | High grade | | Class 2a | 3,000 | 272 | 3,000 | 1,819 |
| 2702 | Male | T1 | Low grade | | Class 2a | 3,000 | 254 | 3,000 | 2,355 |
| 4418 | Male | T1 | High grade | | Class 2a | 3,000 | 195 | 3,000 | 1,461 |
| 2919 | Female | T1 | High grade | | Class 3 | 3,000 | 189 | 3,000 | 2,589 |
| 2004 | Male | T1 | Low grade | | Class 3 | 2,750 | 140 | 2,750 | 1,525 |
| 1670 | Male | T1 | High grade | | Class 2a | 3,000 | 121 | 3,000 | 2,239 |
| 827 | Male | T1 | High grade | | | 3,000 | 110 | 2,706 | 326 |
| 3673 | Male | T1 | High grade | | Class 2a | 3,000 | 55 | 3,000 | 1,978 |
| 2755 | Male | T2-4 | High grade | LumP | | 3,000 | 631 | 3,000 | 1,782 |
| 2815 | Male | T2-4 | High grade | LumU | | 2,500 | 414 | 2,500 | 988 |
| 2368 | Male | T2-4 | High grade | Ba/Sq | | 1,250 | 380 | 1,250 | 246 |
| 1468 | Male | T2-4 | High grade | LumP | | 3,000 | 339 | 3,000 | 1,220 |
| 3472 | Male | T2-4 | High grade | LumP | | 750 | 338 | 750 | 181 |
| 4172 | Male | T2-4 | High grade | LumU | | 750 | 302 | 750 | 112 |
| 1648 | Male | T2-4 | High grade | Stroma-rich | | 2,000 | 278 | 1,896 | 221 |
| 4735 | Male | T2-4 | High grade | | | 1,000 | 239 | 1,000 | 584 |
| 2704 | Male | T2-4 | High grade | NE-like | | 3,000 | 210 | 3,000 | 1,184 |
| 3143 | Male | T2-4 | High grade | LumU | | 3,000 | 209 | 3,000 | 1,149 |
| 3612 | Male | T2-4 | High grade | | | 3,000 | 187 | 2,988 | 313 |
| 2685 | Male | T2-4 | High grade | LumU | | 3,000 | 182 | 3,000 | 533 |
| 1428 | Male | T2-4 | High grade | LumP | | 3,000 | 181 | 2,000 | 968 |
| 4528 | Male | T2-4 | High grade | Stroma-rich | | 3,000 | 159 | 2,688 | 327 |
| 4261 | Female | T2-4 | High grade | Ba/Sq | | 3,000 | 152 | 2,866 | 172 |
| 2514 | Female | T2-4 | High grade | LumP | | 2,500 | 144 | 2,500 | 1,794 |
| 3923 | Female | T2-4 | High grade | LumP | | 3,000 | 130 | 3,000 | 686 |
| 3232 | Male | T2-4 | High grade | LumP | | 3,000 | 116 | 2,995 | 779 |
| 3096 | Male | T2-4 | High grade | Ba/Sq | | 2,750 | 100 | 2,750 | 839 |
| 2868 | Male | T2-4 | High grade | Ba/Sq | | 3,000 | 98 | 2,410 | 139 |
| 3710 | Male | T2-4 | High grade | Ba/Sq | | 3,000 | 90 | 3,000 | 105 |
| 4403 | Male | T2-4 | High grade | Ba/Sq | | 3,000 | 88 | 2,019 | 159 |
| 3170 | Male | T2-4 | High grade | LumU | | 1,750 | 70 | 815 | 93 |
| 2269 | Female | T2-4 | High grade | LumP | | 1,250 | 52 | 586 | 44 |
| 4503 | Female | T2-4 | High grade | Ba/Sq | | 2,250 | 41 | 642 | 107 |

Supplementary Table S2: Univariate- and multivariable cox regression analysis

| Progression-free survival (60 months follow-up) | | |
|---|---|---|
| | HR (95% CI) | *p*-value |
| Univariate analysis | | |
| Class 2a weight (high vs low; n=136, 25 events) | 4.20 (1.75-10.07) | 0.0013 |
| EORTC risk score (>6 vs ≤6; n=136, 25 events) | 14.80 (2.00-109.4) | 0.0083 |
| EAU risk score (high vs intermediate/low; n=134, 25 events) | 9.26 (1.25-68.48) | 0.029 |
| Multivariable model 1 (n=136, 25 events) | | |
| Class 2a weight (high vs low) | 2.58 (1.06-6.29) | 0.036 |
| EORTC risk score (>6 vs ≤6) | 10.07 (1.31-77.08) | 0.026 |
| Multivariable model 2 (n=134, 25 events) | | |
| Class 2a weight (high vs low) | 2.97 (1.22-7.23) | 0.017 |
| EAU risk score (high vs intermediate/low) | 5.92 (0.77-45.45) | 0.087 |

HR = hazard ratio; CI = confidence interval; EORTC = European Organisation for Research and Treatment of Cancer; EAU = European Association of Urology.
